# Activation and inhibition of nonsense-mediated mRNA decay controls the abundance of alternative polyadenylation products

**DOI:** 10.1101/794719

**Authors:** Aparna Kishor, Sarah E. Fritz, Nazmul Haque, Zhiyun Ge, Wenjing Yang, Jun Zhu, J. Robert Hogg

## Abstract

Alternative polyadenylation (APA) produces transcript 3’ untranslated regions (3’UTRs) with distinct sequences, lengths, stability, and functions. We show here that APA products include a class of cryptic nonsense-mediated mRNA decay (NMD) substrates with extended 3’UTRs that gene- or transcript-level analyses of NMD often fail to detect. Transcriptome-wide, the core NMD factor UPF1 preferentially recognizes long 3’UTR products of APA, leading to their systematic downregulation. Further, we find that many APA events consistently observed in multiple tumor types are controlled by NMD. Additionally, PTBP1, previously implicated in direct modulation of co-transcriptional polyA site choice, regulates the balance of short and long 3’UTR isoforms by inhibiting NMD. Our data suggest that PTBP1 binding near polyA sites can drive production of long 3’UTR APA products in the nucleus and/or protect them from decay in the cytoplasm. Together, our findings reveal a widespread role for NMD in shaping the outcomes of APA.

## INTRODUCTION

The 3’ untranslated regions (3’UTRs) of messenger RNAs (mRNAs) coordinate post-transcriptional regulatory mechanisms. The sequence, structure, and length of 3’UTRs determine their interactions with trans-acting factors, thereby controlling transcript functions, localization, and stability (Mayr, 2016). Correspondingly, cells exploit alternative pre-mRNA cleavage and polyadenylation (APA) to generate transcript isoforms with functionally distinct 3’UTRs. A major class of APA events involves differential recognition of multiple potential pA sites within a single last exon (frequently termed tandem APA) (Edwalds-Gilbert et al., 1997). Tandem APA typically produces mRNAs with identical coding sequences but 3’UTRs of different sequence and length.

Changes in the cellular abundance of core cleavage and polyadenylation factors, often observed in cellular differentiation and transformation, result in systematic expression of mRNAs with either short (increased use of stop codon-proximal pA sites) or long (increased use of distal pA sites) 3’UTRs (Gruber et al., 2014; Ji et al., 2009; Kim Guisbert et al., 2006; Lackford et al., 2014; Martin et al., 2012; Shepard et al., 2011; Takagaki and Manley, 1998; Takagaki et al., 1996; Xia et al., 2014; Yao et al., 2012). Along with systemic effects driven by core factor abundance, APA is modulated by RBPs that prevent recognition of potential polyadenylation sites or enhance recruitment of cleavage and polyadenylation factors (Batra et al., 2014; Castelo-Branco et al., 2004; Kyburz et al., 2006; Millevoi et al., 2009; Nazim et al., 2017). While APA has primarily been studied with respect to the regulation of pA site choice in the nucleus, the abundance of APA transcript isoforms may also be controlled at the level of mRNA stability. For example, long 3’UTR products of tandem APA are frequent targets of miRNA-mediated repression, enabled by increased frequency of miRNA binding sites in distal 3’UTR segments (Mayr and Bartel, 2009).

In addition to post-transcriptional control of APA products by miRNAs, the nonsense-mediated mRNA decay pathway (NMD) can regulate the abundance of long 3’UTRs produced by APA (Bao et al., 2016). NMD is a translation-dependent mRNA decay pathway dually responsible for performing RNA quality control and regulation of many apparently normal genes (Kishor et al., 2019a). The NMD pathway is strongly activated by the presence of an exon junction complex (EJC) more than 50 nt downstream of a termination codon (Chamieh et al., 2008; Le Hir et al., 2001; Nagy and Maquat, 1998), but a second major class of NMD targets are mRNAs containing long 3’UTRs (Eberle et al., 2008; Singh et al., 2008). 3’UTR length-sensing by NMD is mediated through UPF1, a highly-conserved RNA helicase that extensively binds mRNA in a sequence-independent fashion due to contacts on the ribose backbone (Chakrabarti et al., 2011; Hogg and Goff, 2010). Increased UPF1 occupancy on an mRNA increases the chances of UPF1 phosphorylation by SMG1, the principal UPF1 kinase (Durand et al., 2016; Kashima et al., 2006). UPF1 phosphorylation is an important signal that favors transcript decay, leading to recruitment of the SMG6 endonuclease and the SMG5/7 complex, which links NMD to decapping and deadenylation enzymes (Eberle et al., 2009; Huntzinger et al., 2008; Loh et al., 2013; Nicholson et al., 2018). Proximal pA site usage may produce short 3’UTRs immune to NMD, while distal pA sites may produce NMD-sensitive mRNAs. However, despite the characterized ability of NMD to recognize and degrade long 3’UTRs, the role of the pathway in post-transcriptional regulation of APA events has not been extensively explored.

The NMD sensitivity of transcripts with long 3’UTRs can be modulated by two cellular proteins that act as protective factors, PTBP1 and hnRNP L (Ge et al., 2016; Kishor et al., 2019b). When these proteins bind near the termination codon (TC) of a mature transcript, they prevent UPF1 accumulation on the 3’UTR and inhibit decay. In addition to their anti-NMD functions, both PTBP1 and hnRNP L have roles in regulation of splicing as well as pA site selection (Attig et al., 2018; Castelo-Branco et al., 2004; Gruber et al., 2018; Huang et al., 2012; Le Sommer et al., 2005). Thus, these proteins may affect the abundance of mRNA isoforms both by regulating their production in the nucleus and by modulating their stability in the cytoplasm.

In the present study, we investigate whether NMD influences the relative abundance of long and short transcript isoforms produced by tandem APA. We find that NMD systematically targets long 3’UTR products of APA due to direct binding of UPF1 to extended 3’UTR segments. Our analyses indicate that many APA products regulated by NMD may not be normally detected by transcript-level approaches, instead requiring interrogation of the relative abundances of individual transcript segments. Further, we find that many instances of apparent APA regulation by PTBP1 can be explained by its role in protection of mRNAs from NMD rather than direct modulation of co-transcriptional pA site selection. Together, our data demonstrate that APA products represent a class of cryptic NMD targets under negative regulation by UPF1 and positive regulation by PTBP1.

## RESULTS

### Analysis of the role of NMD in determining 3’UTR isoform abundance

Many human genes contain multiple potential sites for co-transcriptional cleavage and polyadenylation. Differential selection of polyA sites within the same last exon leads to production of mRNAs with common coding sequences and partially overlapping 3’UTRs. Most transcript-level gene expression analyses include reads that map to mRNA segments shared by multiple isoforms. This means that quantification of long 3’UTR isoforms may be biased by the high abundance of short 3’UTR isoforms that share common 5’UTR and CDS regions, limiting their utility in revealing the impact of NMD on products of APA. To avoid this problem, we chose to pursue a 3’UTR segment-level analysis that considered reads common to multiple transcript isoforms separately from reads that uniquely identify 3’UTR segments generated by differential polyA site choice (Figure 1A).

**Figure 1.**
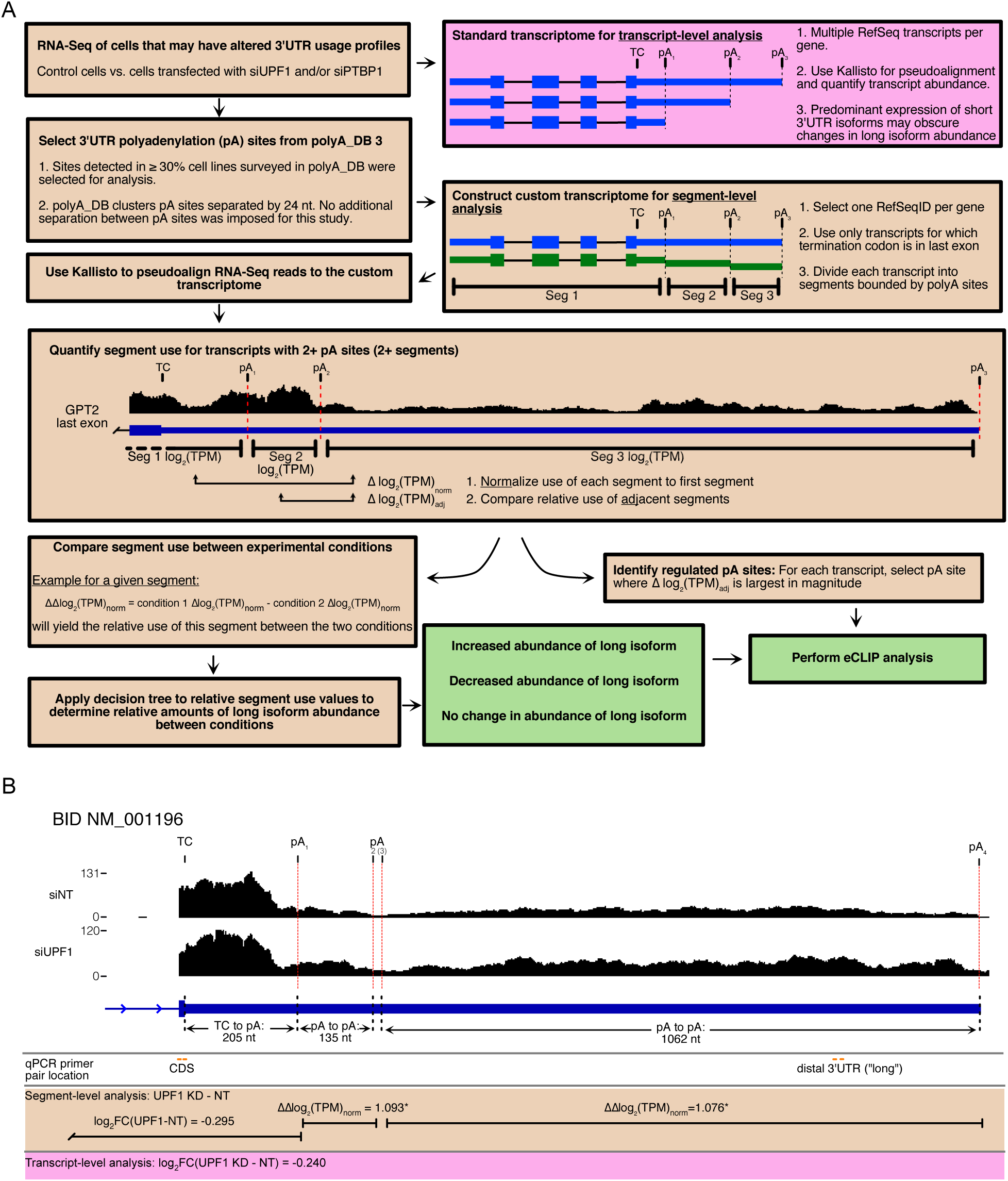
A segment-level RNA-Seq analysis strategy reveals mRNA isoform-specific abundance changes. **(A)** Computational and analytical workflow. See methods for details. **(B)** RNA-Seq trace of the BID locus. Vertical red lines indicate segment boundaries. qPCR primer positions are indicated. Quantification results from segment-level transcript-level analysis are shown. The segment from pA_2_ to pA_3_ was excluded from analysis because it did not meet the TPM cutoff. Stars indicate *P* ≤ 0.05.

To quantify 3’UTR segments potentially regulated by NMD, we first constructed a custom transcriptome based on polyA sites from polyA_DB, a collection of empirically determined polyA sites from multiple cell types (Wang et al., 2018). Specifically, we selected pA sites in 3’UTRs observed in 30% or more of the cell types used to build the polyA_DB (transcripts binned by numbers of polyA sites are shown in Figure S1A). Transcripts were divided into segments based on the locations of these well-supported polyA sites (Figure 1A). The first segment of each transcript used for quantification consisted of the 5’UTR, CDS, and 3’UTR upstream of the first polyA site, followed by separate segments bounded by each additional polyA site. The abundance of each segment was then compared to the abundance of the first segment to determine relative 3’UTR isoform use [Δlog_2_(TPM)_norm_]. To integrate the segment-by-segment analysis into an overall evaluation of changes of isoform abundance in different experimental conditions, we constructed a decision tree (see Methods for details). To determine whether longer or shorter 3’UTR isoforms are favored, the decision tree places the most weight on relative abundance changes of the last 3’UTR segment between conditions, but also accounts for significant abundance changes of internal segments.

We used this strategy first to assess changes in relative transcript segment abundance in HEK-293 cells transfected with non-targeting siRNA or UPF1-specific siRNA to impair NMD (Kishor et al., 2019b). The sensitivity of the segment-level approach is illustrated by transcripts encoding the apoptosis regulator BID (Figure 1B). Segment-level analysis showed that two BID 3’ UTR isoforms were significantly more abundant with siUPF1 than non-targeting siRNA transfection [ΔΔlog_2_(TPM)_norm_ of 1.093 and 1.076, respectively]. In contrast, a transcript-level analysis showed no increase in the abundance of long 3’UTR-containing BID mRNAs between these two conditions because the high abundance of short 3’UTR BID mRNAs masked the change in levels of the less abundant long isoforms. Additional example traces are provided in Figure S1B.

### UPF1 systematically suppresses long 3’UTR products of APA

The segment-level analysis detailed above revealed that UPF1 knockdown (KD) induced a systematic shift toward increased usage of long 3’UTR isoforms (Figures 2A and S2A; Table S1). It should be noted that the results of the decision tree sometimes differ from the comparison of the last segment (Figure 2A, red points below threshold) because a significant abundance change occurred at an internal segment of the 3’UTR rather than the last segment. Of the analyzed genes that underwent significant shifts in the usage of long 3’UTR segments, the vast majority had increased abundance of long isoforms when UPF1 was knocked down (1456 of 5981 total genes analyzed; Figure 2A), suggesting that NMD preferentially clears long 3’UTR transcript isoforms. Notably, transcripts with enhanced long 3’UTR isoform expression upon UPF1 KD did not exhibit increased abundance in a traditional isoform-level RNA-Seq analysis (Figure 1A and 2B) and thus would not be identified as NMD targets.

**Figure 2.**
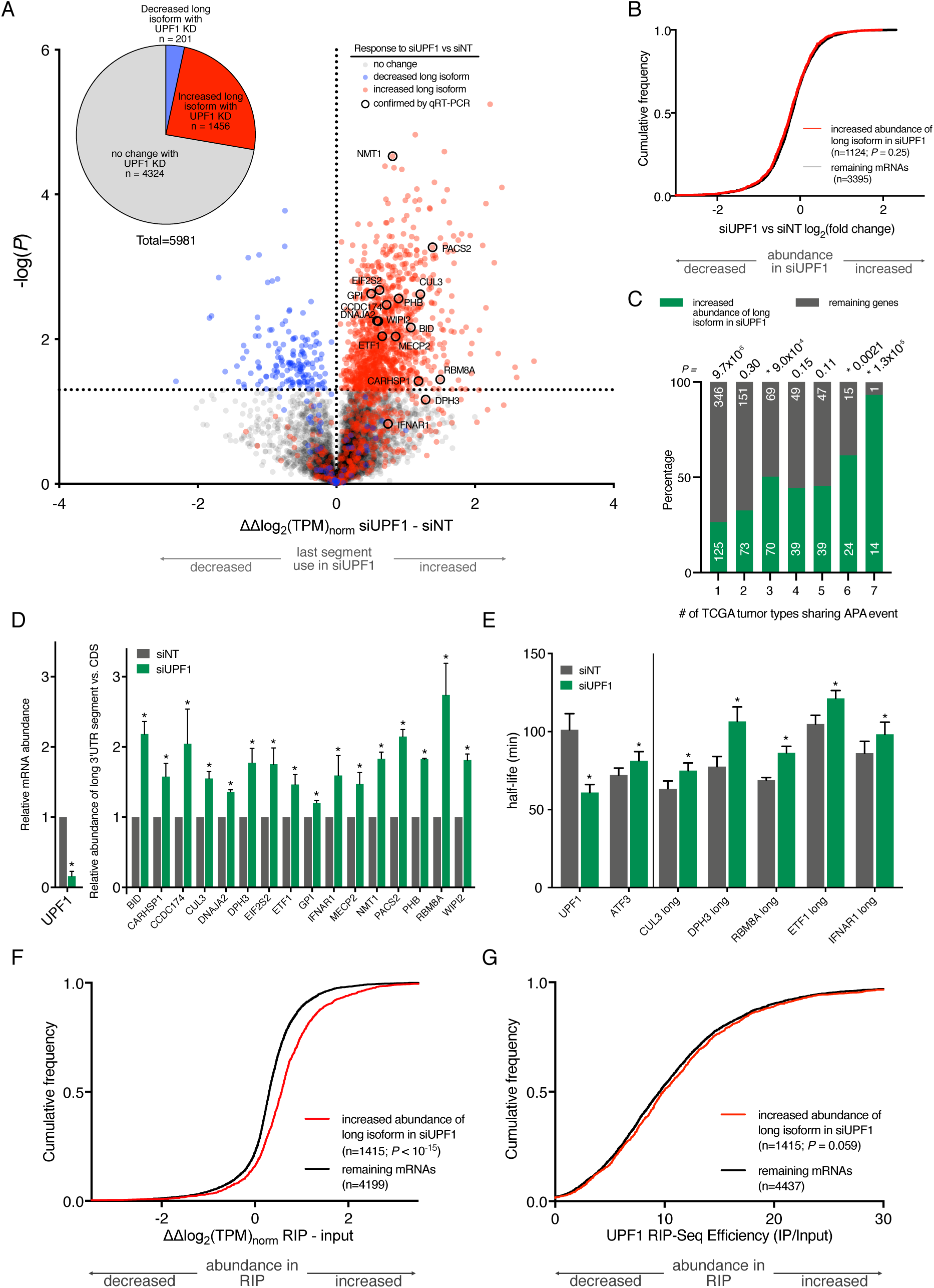
Compromise of NMD globally increases abundance of long 3’UTR isoforms. **(A)** Volcano plot representing ΔΔlog_2_(TPM)norm between non-targeting and UPF1 knockdown of the last segment. Horizontal line indicates the significance threshold *P* ≤ 0.05 (n = 3). Points are color coded in accordance with the results of the decision tree as indicated in the figure key. Inset: results of the decision tree. **(B**) CDF plot of log_2_FC of transcripts grouped by the classes in (A) quantified using transcript-level analysis. Significance from the Kolmogorov-Smirnov (K-S) test. **(C)** Proportion of transcripts with increased long isoforms upon UPF1 knockdown in common with the set of mRNAs identified as APA targets in TCGA (Xia et al., 2014). DaPars mRNAs are binned by the number of tumor types that share the APA event. The significance of the over-or under-representation of these transcript bins in the UPF1 KD set was evaluated by a two-tailed binomial test (\**P*≤0.05 for over-represented bins). **(D)** Relative abundance of long isoforms compared to transcript CDS in siNT and siUPF1 knockdown conditions. siUPF1 values were normalized to the non-targeting condition (n = 3). Significance was determined by two-tailed Student’s t-test (\**P*≤0.05; error bars = 1SD). **(E)** Metabolic labeling to determine the half-lives of the long isoform under conditions of UPF1 knockdown or non-targeting siRNA (n = 4). Genes to the left of the vertical line are controls. Significance was determined by using a two-tailed Student’s t-test (\**P*≤0.05; error bars = 1SD). **(F)** CDF plot of log_2_FC of the transcript abundance in UPF1 affinity purifications compared to the input. Genes were divided into classes from (A) according to their response to siUPF1 knockdown. Significance was calculated by K-S test. **(G)** RIP-Seq data from (D) quantified using transcript-level analysis. Transcripts were grouped according to the segment-level analysis as in (D), and the figure shows the log_2_FC of these classes. Significance was calculated by K-S test.

In order to investigate the relationship between the NMD-sensitive isoforms identified here and APA events in disease, we compared our dataset to a previous analysis that demonstrated pervasive shortening of 3’UTRs in seven tumor types from the Cancer Genome Atlas (TCGA) (Hu et al., 2017; Xia et al., 2014). Of the 1,062 genes that were identified as APA targets in one or more tumor types and met read count criteria in the present study, we found that 384 were present in our set of 1456 genes with increased long 3’UTR isoform abundance in response to UPF1 depletion (Figure 2C). In each of the seven tumor types, our analysis identified a consistent 43-48% of the APA target genes as potential NMD targets (Figure S2B). Interestingly, the overlap between long 3’UTR isoforms repressed by NMD and the previously identified APA events in cancers was greater for events represented in multiple tumor types. Of the 39 genes affected in six tumor types, 24 exhibited increased long 3’UTR isoform expression in UPF1 KD, and 14 of 15 genes identified in all tumor types were identified here as UPF1 targets (Figure 2C and S2C). These data suggest that NMD may be an important unappreciated modifier of 3’UTR isoform expression in many cancers.

We selected several genes with increased relative abundance of long 3’UTR isoforms under conditions of UPF1 KD to validate by comparing amplicons generated from qPCR primers sited in the CDS and 3’UTR extension (Figures 1B and 2D). To distinguish between control of 3’UTR isoform abundance through pA site choice *versus* differential decay by NMD, we used metabolic labeling with 5-ethynyluridine. In this analysis, the long 3’UTR isoform transcripts were generally turned over more rapidly than the overall transcript pool derived from each gene but were stabilized with UPF1 KD, suggesting that the effect of UPF1 on APA product abundance is indeed via RNA decay (Figure 2E and S2D).

### UPF1 preferentially recognizes extended 3’UTR segments

For insight into whether UPF1 directly regulates 3’UTR isoform usage, we analyzed RIP-Seq data of mRNAs associated with affinity purified UPF1 (Table S2). In general, mRNA recovery was robust, but mRNAs containing distal 3’UTR segments were preferentially bound to UPF1 (Figure S2E), resulting in a transcriptome-wide enrichment of the last 3’UTR segment in the RIP compared to the input RNA (Figure S2F). Importantly, transcripts containing distal segments with increased expression upon siRNA KD of UPF1 (from Figure 2A) exhibited significantly increased relative recovery with affinity purified UPF1 (Figure 2F). Enrichment of UPF1-associated RNA *versus* input evaluated without 3’UTR segmentation did not clearly identify this relationship (Figure 2G).

### Multiple roles of PTBP1 in determining 3’UTR isoform abundance

The multifunctional RNA binding protein PTBP1 has been implicated in regulation of pre-mRNA cleavage and polyadenylation (Attig et al., 2018; Castelo-Branco et al., 2004; Le Sommer et al., 2005). We have also shown that PTBP1 inhibits decay of potential NMD target mRNAs (Ge et al., 2016). We therefore investigated how direct PTBP1 regulation of pA site selection might intersect with its function in inhibiting NMD. To begin to deconvolve these effects, we analyzed relative 3’UTR isoform abundance in cells depleted of PTBP1, either alone or together with UPF1 KD.

Analysis of our RNA-Seq data through the decision tree revealed similar numbers of genes undergoing increases (1,561 genes) and decreases (1,677 genes) in relative abundance of long isoforms upon PTBP1 KD compared to non-targeting siRNA (Figure 3A; Table S3), consistent with the notion that PTBP1 can affect mRNA isoform abundance through multiple mechanisms. We elected to focus on transcripts that had decreased long isoform abundance with PTBP1 KD, as these are candidates for PTBP1-mediated protection from NMD (Ge et al., 2016). In the majority of transcripts in this group, expression of the extended 3’UTR isoforms was restored when PTBP1 and UPF1 were simultaneously knocked down as compared to PTBP1 KD alone (n = 1,050) (Figures 3A, 3B, S3A, upper left quadrant, and S3B). This finding indicates that a substantial fraction of PTBP1-dependent changes in 3’UTR isoform abundance may be attributed to reduced NMD efficiency. A subset of genes (n = 627) underwent a decrease in long isoform abundance with PTBP1 KD without restoration by simultaneous UPF1 KD, behavior consistent with either a direct effect of PTBP1 on alternative pA site usage in the nucleus or PTBP1-mediated inhibition of other RNA decay pathways (Figure 3B). We used qPCR to confirm that PTBP1 KD decreased the relative amount of the long isoform of selected transcripts in this class (Figure 3C). Further, decay of long 3’UTR isoforms was enhanced in the absence of PTBP1 and abrogated by UPF1 KD (Figure 3D), an effect obscured in an isoform-independent approach (Figure S3C).

**Figure 3.**
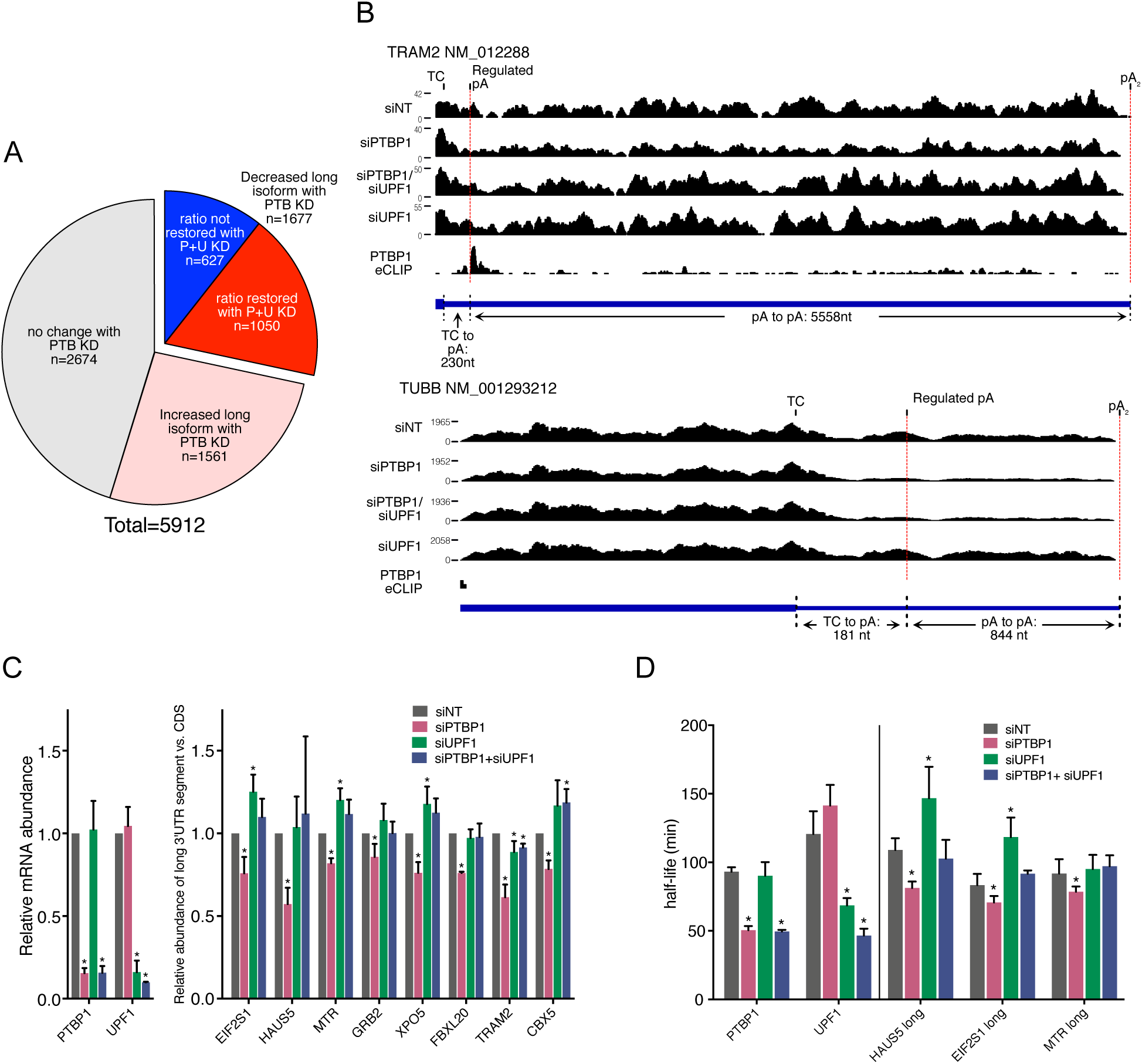
PTB knockdown exposes some long mRNA isoforms to NMD. **(A)** Pie chart representing classes of mRNAs revealed by independent or simultaneous PTBP1 and UPF1 knockdown. The class of transcripts for which PTBP1 knockdown resulted in decreased long isoform abundance is divided into mRNAs that had their relative isoform abundance ratio restored when UPF1 was knocked down simultaneously with PTBP1 and those that did not. **(B)** Representative RNA-Seq traces for transcripts that had their isoform abundance restored with UPF1 + PTBP1 knockdown (TRAM2) and those that did not (TUBB), along with the PTBP1 eCLIP reads. **(C)** Fold change of long isoform expression compared to the CDS of the transcript under the indicated knockdown conditions. Changes were normalized to the non-targeting condition (n = 3). Controls are shown to the left. Significance was determined by using a two-tailed Student’s t-test (\**P* ≤ 0.05; error bars = 1SD). **(D)** Metabolic labeling to determine the half-lives of the long isoform under the indicated conditions (n = 4). Transcripts to the left of the vertical line are controls. Significance was determined byusing a two-tailed Student’s t-test (\**P* ≤ 0.05; error bars = 1SD).

### Distinct PTBP1 binding profiles in NMD inhibition and APA

Binding of PTBP1 in the vicinity of pA sites has supported the hypothesis that it directly regulates APA (Attig et al., 2018; Gruber et al., 2018), while PTBP1 association near TCs correlates with NMD inhibition (Ge et al., 2016). To investigate the relationship between PTBP1 binding and its potential nuclear and cytoplasmic roles in determining APA outcomes, we performed eCLIP (Van Nostrand et al., 2016). In order to analyze PTBP1 binding events of interest, we first identified the pA sites with the biggest abundance changes (marked “regulated pA” in Figure 3B and Figure S3B). Importantly, 50% of the regulated pA sites were within ~500 nt of the termination codon (Figure 4A). Metagene plots of PTBP1 eCLIP peaks near pA sites revealed markedly different PTBP1 binding profiles for the three major gene classes (Figures 3A and 4B), particularly upstream of the regulated pA sites. Further analysis of the genes with decreased long isoform abundance in the absence of PTBP1 revealed PTBP1 binding near regulated pA sites also tends to be close to the TC (Figure 4C). When this population was subdivided by response to the double KD (Figure 4D), NMD-dependent events were distinguished by a shift in PTBP1 binding enrichment toward the stop codon (green trace), while NMD-independent events were marked by PTBP1 peaks closer to the pA site (red trace). We propose that the broad enrichment of PTBP1 upstream of pA sites is at least partially a consequence of PTBP1 bound near the TC to inhibit NMD.

**Figure 4.**
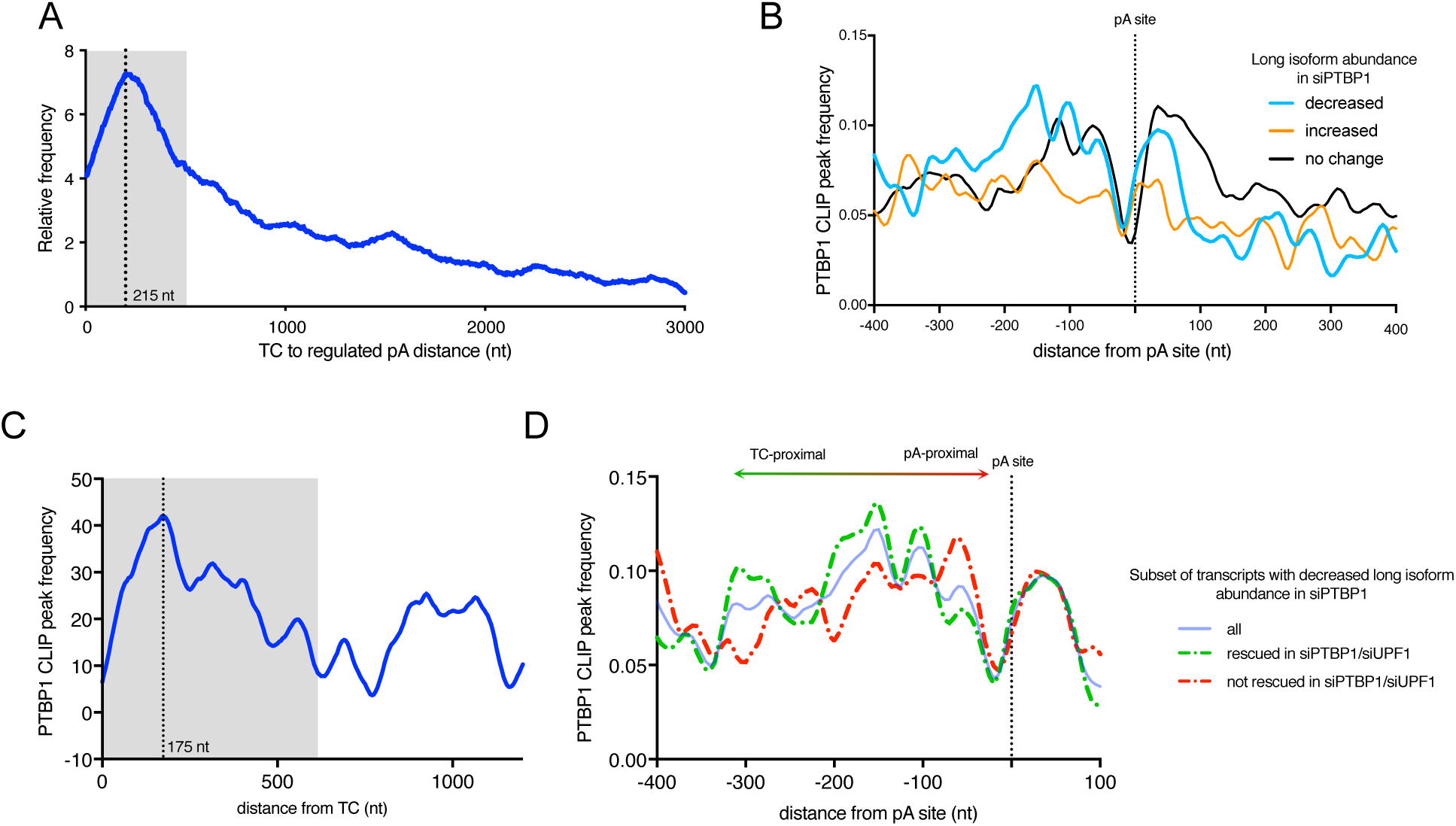
PTBP1 binding near pA sites can be associated with NMD inhibition. **(A)** Histogram of the distance between the regulated pA sites and the termination codon. Grey shaded region represents 50% of the area under the curve. **(B)** eCLIP metagene plot for PTBP1 binding sites near pA sites that define segments with the greatest abundance change. Transcripts are divided into classes based on results of segment-level analysis. **(C)** Histogram of distance of PTBP1 eCLIP peaks from termination codons at regulated pA sites for transcripts that exhibited decreased abundance of the long isoform with PTBP1 KD. Grey shaded region represents 50% of the area under the curve. **(D)** eCLIP metagene plot for PTBP1 binding near regulated pA sites for the transcripts with decreased long isoform abundance with PTBP1 knockdown, subdivided by whether long isoform abundance was restored with simultaneous UPF1 knockdown.

## DISCUSSION

While co-transcriptional selection of pA sites is critical for generating APA isoforms, the abundance of those isoforms is also subject to regulation at the level of RNA stability. Here, we uncover a widespread role for NMD in regulating long 3’UTR products of APA (Figure 1). Supporting a direct role for UPF1 in this process, we show that mRNAs containing extended 3’UTR isoforms are preferentially recovered in UPF1 RIP-Seq and that long 3’UTR APA products of selected genes are destabilized in the presence of UPF1 (Figure 2). Further, we present evidence that the activity of NMD in suppressing expression of long 3’UTR products of APA is counteracted by the protective protein PTBP1. Consistent with previous reports, our data implicate PTBP1 both in enhancement and repression of APA (Attig et al., 2018; Castelo-Branco et al., 2004; Gruber et al., 2018; Le Sommer et al., 2005). Importantly, however, our data suggest that a major way that PTBP1 determines the outcomes of APA is by protecting long 3’UTRs from NMD (Figure 3). Transcriptome-wide analyses show that PTBP1 binding near pA sites is also frequently in close proximity to the TC, raising the possibility that recruitment of PTBP1 to pA sites can affect cleavage site selection, NMD susceptibility, or both (Figure 4).

Our discovery of over a thousand genes that show evidence of UPF1-mediated control of APA product abundance illustrates that prior approaches substantially underestimate the scope of long 3’UTR turnover by NMD. NMD target isoforms are often expressed at lower levels than non-NMD target isoforms from the same locus, meaning that non-target mRNAs can obscure the effects of NMD if gene- or transcript-level analyses are used. It has long been recognized that examination of the use of individual exons is necessary to understand the interplay between alternative splicing and NMD (McGlincy and Smith, 2008). Our data show that a similar approach is needed to uncover the role of NMD in determining the outcomes of APA. This approach may be especially important when considering results generated from standard laboratory lines such as HEK-293 or other transformed cells. These cells tend to have enhanced use of proximal poly-A sites, resulting in a low basal level of long 3’UTR isoform expression (Mayr and Bartel, 2009). Thus, any NMD-induced changes in the abundance of the long isoforms will be hidden unless analysis pipelines are used that are sensitive to this reservoir of cryptic NMD targets.

PTBP1 (or other RBP) binding near pA sites is used to support their identification as direct regulators of APA (Attig et al., 2018; Gruber et al., 2018). In the case of PTBP1, its role has been confirmed by previous biochemical studies that found that PTBP1 can directly affect pA site choice in the nucleus by enhancing or inhibiting recognition of pA sites (Castelo-Branco et al., 2004; Le Sommer et al., 2005). We find evidence of a unique pattern of PTBP1 binding near many pA sites upstream of PTBP1-dependent segments that are rescued by NMD inhibition (Figure 4D), suggesting that these PTBP1 binding events actually regulate NMD sensitivity by virtue of their increased proximity to the TC. Together, these data show that binding of *trans*-acting factors cannot be assumed to be evidence for a direct effect on nuclear pA site selection, as they may represent binding events required to maintain cytoplasmic stability of the mRNA.

The concept that alternative splicing can generate NMD substrates as a regulatory mechanism is well established, but the coupling of APA to NMD to achieve regulation has not been similarly explored. We validated several genes involved in translation and decay as undergoing UPF1-dependent changes in 3’UTR isoform abundance (Figure 2D). In addition to these targets, we also found that the long 3’UTR isoform of MeCP2 is repressed by UPF1 (Figure S1B). MeCP2 is mutated in the neurodevelopmental disorder Rett syndrome (Lyst and Bird, 2015), and alteration of MeCP2 APA by NUDT21 copy number variations has been implicated in neuropsychiatric disease (Gennarino et al., 2015).

Finally, we find a surprising degree of overlap between UPF1 targets and genes previously found to undergo 3’UTR “shortening” in cancer. Multiple lines of evidence suggest that preferential use of proximal pA sites in cancer is associated with increased expression of core cleavage and polyadenylation proteins (Turner et al., 2018), but our data indicate that nuclear control of 3’UTR isoform expression in cancer may be systematically augmented by NMD. These findings raise two possibilities that merit further investigation: first, NMD efficiency may contribute to 3’UTR isoform expression changes in cancer and second, APA may be in part used by cancer cells to evade NMD.

## ACKNOWLEDGEMENTS

We thank Joseph Chapman for critical evaluation of the manuscript, Fayaz Seifuddin for discussion of the kallisto analysis, Thomas Baird for metabolic labeling samples, and the NHLBI DNA Sequencing and Genomics Core. This work was supported by the Intramural Research Program, National Institutes of Health, National Heart, Lung, and Blood Institute and utilized the computational resources of the NIH HPC Biowulf cluster. (http://hpc.nih.gov).

## AUTHOR CONTRIBUTIONS

AK: experimental framework, steady state qPCR, metabolic labeling and related qPCR, RNA-Seq data analysis and pA analysis decision tree development, manuscript; SEF: RIP-Seq; NH: PTBP1 eCLIP; ZG: samples for the RNA-Seq library for the KD experiments; WY: RNA-Seq analysis; JZ: experimental framework; JRH: experimental framework, RNA-Seq data analysis and pA analysis decision tree development, eCLIP analysis, manuscript.

## DECLARATION OF INTEREST

The authors declare no competing interests.

## METHODS

### Cultured cells and siRNA

HEK-293 Tet-Off cells and Flp-In T-REx-293 cells (ThermoFisher Scientific) at 37°C in ambient oxygen and 5% CO_2_ in DMEM (Gibco #11965-092) supplemented with 10% fetal bovine serum (Gibco) and a 1% penicillin, streptomycin, and L-glutamine mixture (Gibco). Human Flp-In T-REx-293 cells expressing 3xFLAG-PTBP1 or CLIP-UPF1 were generated following the manufacturer’s protocol (ThermoFisher Scientific). In brief, Flp-In T-REx-293 cells were transfected with pcDNA5/FRT/TO-PTBP1 or pcDNA5/FRT/TO-CLIP-UPF1 and pOG44 plasmids (ThermoFisher Scientific) with TurboFect or LipoFectamine 3000 transfection reagents according to the manufacturers’ suggestions (ThermoFisher Scientific). Polyclonal cells were treated with doxycyline hyclate (1 μg/mL final concentration, ThermoFisher Scientific) for 48-72 hours for induction of transgene expression, and transgene expression was confirmed by western-blot analysis (anti-FLAG, Sigma; anti-UPF1, Bethyl Laboratories).

siRNAs used were non-targeting siRNA: Silencer Select negative control #2 siRNA (ThermoFisher Scientific #4390846); UPF1 siRNA: GAUGCAGUUCCGCUCCAUUUU; PTPB1 siRNA: CUUCCAUCAUUCCAGAGAAUU (ThermoFisher Scientific; Mendell *et al*, 2004).

### Plasmids

Full-length PTBP1 cDNA was amplified by RT-PCR from HEK-293 total RNA and cloned into the tetracycline-inducible expression vector pcDNA5/FRT/TO (ThermoFisher Scientific), which was modified to harbor an N-terminal 3XFLAG tag (Haque et al., 2018).

### RNA-Seq

The RNA-Seq datasets used in this study reflecting the transcriptomes of cells with UPF1 KD, PTBP1 KD and dual UPF1+PTBP1 KD have been documented in our previous studies (Kishor et al 2019, Ge et al 2016). Sequencing data are available from the NCBI GEO database.

Accession GSE109143 (3 trials for siNS and 3 trials for siUPF1) (Kishor et al 2019) and GSE59884 (2 trials each for 4 PTBP1 experimental conditions) (Ge et al 2016).

### PTBP1 eCLIP

eCLIP was performed as previously reported by Nostrand et al., 2016, with the following modifications. By using click-chemistry conjugation, 3’-adapters were non-radioactively labeled with infrared (IR) dye at the 3’ end. Labelling and purification of oligonucleotides was performed essentially as described by Zarnegar et al., 2016. In brief, modified adapters with an azide group at the 3’ end (Modified-RNA_X1A: /5Phos/rArUrArUrArGrG rNrNrNrNrNrArGrArUrCrGrGrArArGrArGrCrGrUrCrGrUr GrUrArG/3AzideN/; Modified-RNA_X1B: /5Phos/rArArUrArGrCrArNrNrNrNrNrArGr ArUrCrGrGrArArGrArGrCrGrUrCrGrUrGrUrArG/3AzideN/) were purchased from IDT and resuspended at 100 μM in water. 45 μL adapter was mixed with 5 μL 10 mM IRdye-800CW-DBCO (LiCor) and incubated for 2 hr at 37°C. Reactions (50 μL) were mixed with 1200 μL buffer PNI (Qiagen buffer PB with 1.5 volumes 100% isopropanol) and purified with five QIAquick nucleotide removal columns (Qiagen). 250 μL aliquots of the reaction were dispensed in each column and centrifuged at 6000 rpm for 30 seconds. Columns were washed with 0.75 mL 80% ethanol and dried for 2 min by centrifugation at 13,000 rpm. Labeled oligos were eluted from each column in 50 μL water by centrifugation for 1 min at 13,000 rpm. Labelling efficiency was assessed by imaging dot blots on an Odyssey Imaging System (LI-COR Biosciences).

For each eCLIP experiment, 2×10^7^ cells were lysed on ice for 15 min in 1 mL lysis buffer containing 50 mM Tris-HCl pH 7.4, 100 mM NaCl, 1% NP-40, 0.1% SDS, 0.5% sodium deoxycholate, Halt protease inhibitor cocktail (ThermoFisher Scientific). Cells were passed through 27-gauge needles several times before sonication (Biorupter, Diagenode) for 3 min at low setting with 30 second on/30 second off duty cycle at 4°C. A mix of 10 μL1:25 diluted RNase I (Thermo Scientific) and 5 μL Turbo DNase (ThermoFisher Scientific) was added, and lysates were immediately transferred to a Thermomixer (Eppendorf) for incubation for 5 min at 37°C with mixing at 1200 rpm. Next, 10 μL RNase inhibitor (ThermoFisher Scientific) was added, and tubes were immediately transferred to ice. Lysates were cleared by centrifuging at 15,000*g* for 15 minutes at 4°C and transferred to new tubes; 50 μL was saved for RNA and protein analysis. 10 μg Ab coupled with 100 μl of proteinA/G magnetic beads (Pierce) was added to each lysate and mixed for 2 hours at 4°C. IP samples were washed twice with high salt wash buffer (50 mM Tris-HCl pH 7.4, 1 M NaCl, 1mM EDTA, 1% NP40, 0.1% SDS, 0.5% sodium deoxycholate), once with wash buffer (20 mM Tris-HCl pH 7.4, 10 mM MgCl_2_), and twice with FastAP buffer (10 mM Tris pH 7.5, 5 mM MgCl_2_, 100 mM KCl, 0.02% Triton X-100).

Beads were treated first with alkaline phosphatase (8 μL FastAP, ThermoFisher Scientific) for 15 min and then with 7 μL polynucleotide kinase (ThermoFisher Scientific) for 20 min at 37°C. Beads were washed once with wash buffer, twice with high-salt wash buffer, twice with wash buffer, and finally twice with ligase buffer (50 mM Tris-HCl pH 7.5, 10 mM MgCl_2_). 2.5 μL of each IR-labeled adapter was ligated to RNAs bound to beads. Beads were washed consecutively with wash buffer, high-salt wash buffer, and again with wash buffer. Finally, RNA-protein complexes were eluted from the beads in 1X NuPAGE LDS buffer (ThermoFisher Scientific) containing 0.1 M DTT at 70°C for 10 min.

Immunopurified and input materials were run on a 4-12% Bis-Tris protein gel (ThermoFisher Scientific), and protein-linked RNA was transferred to a nitrocellulose membrane (iBlot, ThermoFisher Scientific). Membranes were scanned on an Odyssey Imaging System (LI-COR) or Typhoon Gel and Blot Imaging System (GE Healthcare), and segments of membrane corresponding to the protein band of interest and ~75 kD above were excised and cut into small pieces. Size-matched regions of nitrocellulose membrane from input samples were also excised for RNA extraction. 200 μL proteinase K solution containing 100 mM Tris pH 7.5, 50 mM NaCl, 1 mM EDTA, 0.2% SDS, and 10 μL proteinase K (0.8 U/μL, NEB) was added and mixed for 60 min at 50°C in a Thermomixer. 200 μl saturated phenol-chloroform, pH 6.7, was added and mixed at 1400 rpm for 10 min at 37°C in a Thermomixer. Tubes were centrifuged for 30 seconds at 13,000 rpm, and liquid containing eluted RNA fragments was transferred to 2 mL Heavy Phase Lock Gel tubes (5Prime), which were agitated for 5 minutes at 1200 rpm at 37°C. Tubes were centrifuged for 5 minutes at 13,000 rpm, and 1 mL of chloroform was added and mixed by inverting several times, followed by centrifugation for 5 min at 13,000. Aqueous phases were carefully transferred to new tubes, and RNA was purified using Zymo RNA Clean & Concentrator-5 columns (Zymo Research). RNA was eluted in 10 μL water.

Input RNAs eluted in 10 μL ddH_2_O were first treated with FastAP (2.5 μL FastAP buffer, 0.5 μL RNase inhibitor, and 2.5 μL FastAP enzyme) for 15 min at 37°C. Input RNAs were then mixed with 45 μL ddH_2_O, 20 μL 5XPNK buffer pH 6.5, 1 μL 0.1 mM DTT, 1 μL Turbo DNase (ThermoFisher Scientific), 7 μL T4 Polynucleotide Kinase (NEB) for 15 min at 37°C. Input RNAs were then cleaned up with 20 μL Dynabeads MyONE Silane (ThermoFisher Scientific) and eluted in 10 μL ddH_2_O, 5 μL of which was then used for 3’-linker (RiL19) ligation. First, 5 μL input RNA was mixed with 1.5 μL 100% DMSO and 0.5 μL 40 μM RiL19, incubated at 65°C for 2 min, and immediately transferred on ice. Ligation mix containing 2 μL 10X NEB Ligase buffer, 0.2 μL 1 M ATP, 0.2 μL RNase inhibitor, 0.3 μL 100% DMSO, 8 μL 50% PEG 8000. 1.3 μL RNA ligase (NEB), and 1.5 μL ddH_2_O was added to each input RNA mix and incubated at 27°C for 2 hr with gentle agitation every 15 min. Input RNA samples were then cleaned up with 20 μL Dynabeads MyONE Silane (ThermoScientific) and eluted in 10 μL ddH_2_O. Input and immunopurified RNA samples were used for reverse transcription using SuperScript IV (ThermoScientific). First, 0.5 μL RT primer (AR17, 20 μM) was mixed with 10 μL RNA, heated on 65°C for 4 minute and immediately transferred on ice.

Reverse transcriptase reaction mix containing 1.5 μL ddH_2_O, 1 μL dNTPs (10 mM), 4 μL 5X reaction buffer, 1 μL 0.1 M DTT, 1 μL RNase Inhibitor (ThermoFisher Scientific), 1 μL SuperScript IV was added, followed by incubation for 45 min at 55°C and 30 min at 62°C. Excess primer and free nucleotides were removed from the cDNA with ExoSAP-IT kit (ThermoFisher Scientific). 3.5 μL ExoSAP-IT mix was added to each sample and kept at 37°C for 15 min, after which 1 μL 0.5 M EDTA was added. Subsequently, RNA was removed from the cDNA by adding 3 μL 1M NaOH and incubating at 70°C for 12 min. Reactions were neutralized by adding 3 μL 1M HCl. cDNAs were purified with 10 μL MyONE Silane beads and eluted in 5 μL of 5 mM Tris-HCl pH 7.5. The 5’ adapter (3Tr3) was ligated (0.8 μL 80 μM stock, 2.0 μL RNA ligase buffer, 0.1 M ATP, 9.0 μL 50% PEG-8000, 1.5 μL RNA ligase, 1.1 μL H_2_O) to the 5’-end of cDNAs on bead, and cDNAs were further cleaned up with MyONE Silane beads. cDNAs were eluted and quantified by qPCR by using 1 μL 1:10 diluted cDNA as a template, and finally 12.5 μL cDNA was PCR amplified using Accuprimer Pfx SuperMix (ThermoFisher Scientific). PCR products were purified with AmpureXP beads (Beckman Coulter) and run on 3% low-melt agarose gels. DNA was extracted from gel slices corresponding to 175-350 bp using Qiagen MinElute columns (Qiagen). Purified cDNA libraries were eluted in 12.5 μL buffer EB, quantified in a Qubit Fluorometer (ThermoFisher Scientific), and sequenced on an Illumina HiSeq 3000.

### eCLIP data analysis

RNA-Seq data from eCLIP IP and input samples was analyzed using the ENCODE eCLIP-seq Processing Pipeline (Van Nostrand et al., 2018). Raw fastq files were trimmed with Cutadapt as described (Martin, 2011), followed by removal of sequences derived from repetitive elements with STAR aligner, using human RepBase sequences (Bao et al., 2015; Dobin et al., 2013). Remaining reads were mapped to the hg19 human genome release with STAR aligner. PTBP1 eCLIP peaks were called with Clipper software and normalized vs the size-matched input control as described (Van Nostrand et al., 2016). For metagene analysis, the pA sites marking the largest fold-change in expression of adjacent 3’UTR segments consistent with the decision tree evaluation were used as “regulated pA sites.” As controls, random pA positions from non-regulated genes were selected. MetaPlotR was used to determine peak density at the indicated positions; histograms in Figure 4 were generated by second-polynomial smoothing (Prism software) (Olarerin-George and Jaffrey, 2017).

### Segment-level analysis and decision tree design

#### Creation of the custom transcriptome for polyadenylation analysis

The analysis of the NMD RNA-Seq datasets we use in this study is designed to exploit the pA sites that have been identified through other work. We elected to use the PolyA_DB 3 repository for the human transcriptome curated by Bin Tian’s group (Wang et al., 2018). Our interest is in tandem 3’ UTRs, so we extracted the pA sites annotated as “3’ UTR” for each transcript from the database. We then associated a RefSeq ID to each transcript: for each gene that has multiple mRNA isoforms, the isoform that covered the most 3’ UTR pA annotations was selected. Additionally, we excluded genes in which the termination codon was not in the last exon. This set includes 15,844 transcripts. Genes not represented in PolyA_DB 3 were not evaluated. Once the transcripts had RefSeq IDs assigned, we excluded pA sites that appeared in less than 30% of datasets referenced in PolyA_DB 3. In order to create the custom transcriptome, each transcript was sectioned based on the pA sites. The first section of each transcript runs from the annotated 5’ end of the isoform to the first pA site from the database. Each subsequent section is from the previous pA site to the next pA site, including pA sites beyond the annotated 3’ end of the transcript (schematized in Figure 1A). To summarize, transcripts were divided into as many segments as they have commonly-used pA sites (e.g., if a transcript has 3 pA sites it will be divided into 3 segments: the first from annotated 5’ end to first pA site, the second from first pA site to second pA site, and the third from second pA to third). Each of these sections become independent units for quantification.

#### Pseudoalignment and quantification for polyadenylation analysis

In preparation for pseudoalignment using kallisto (Bray et al., 2016), the coordinates of the custom transcriptome were used to create a FASTA file of the sequence for each section. The index file for kallisto alignment was created using default the k-mer size of 31 and the “make-unique” flag. The reads from our RNA-Seq datasets were pseudoaligned to this index file using the bias, rf-stranded, and bootstrap = 1000 settings. From the resulting data, segments with mean TPMs in the non-targeting condition of less than 0.5 were excluded from analysis.

In the following explanation, “experimental condition” signifies the KD experiment that was evaluated by RNA-Seq (for instance UPF1 KD). “Trial” signifies a single replicate (for instance there are 3 trials for the UPF1 KD experimental condition). All calculations and scripting were done in jsl for the JMP software package. For this study, abundance changes were quantified in 3 ways in order to obtain pA isoform-specific information. First, the log_2_FC of the first segment of each transcript between experimental conditions (for instance non-targeting vs UPF1 KD) was calculated as well as its P value using a Student’s two-tailed t-test (for PTB datasets, a cutoff of 0.2 was used instead of a P value). Second, for each trial, the log_2_FC of every segment compared to the first segment was calculated (this is the normalized difference, or Δlog_2_(TPM)_norm_ for each segment as indicated in Figure 1A). The P value for ΔΔlog_2_(TPM)_norm_ between conditions were calculated (for instance, the difference in Δlog_2_(TPM)_norm_ from each of the 3 nontargeting trials and the 3 UPF1 KD trials). This comparison results in normalization and comparison between conditions of the read density for each segment of each transcript to the density of the first segment. The third quantification is a measurement of the change in use of adjacent segments between conditions. In order to generate this comparison, the log_2_TPM of each segment was compared to the segment immediately prior to it in each trial (this is the difference in segment use or Δlog_2_(TPM)_adj_ for each segment as indicated in Figure 1A). Significance and comparison calculations between conditions was performed as above. At the end of this calculation, the significance and difference of adjacent segment use between conditions were calculated. These changes and their significance were used in the decision tree that evaluates whether longer 3’UTR isoforms are more used in specific experimental conditions than others (see below).

#### Decision tree design

The analytical strategy of the decision tree was based on the quantification of the segment-by-segment changes which were then converted to an overall score to determine whether the longer 3’UTR isoforms were more abundant. The position of the segment in the 3’ UTR is important. In the case of the first segment (which includes the 5’ UTR and the CDS), if the change of the Δlog_2_(TPM) was not significant between conditions, the abundance of that segment between conditions was considered identical. If the change of the Δlog_2_(TPM) was significant between conditions, the abundance of that segment between conditions was marked as higher in one condition over the other. For internal 3’ UTR segments (for instance, segments 2, 3, and 4 of a 5-segment transcript), if the P value of the change in Δlog_2_(TPM)_norm_ between conditions (ΔΔlog_2_(TPM)_norm_) was not significant, then the segments were considered to be used equally between the conditions. If this value was significant and the ΔΔlog_2_(TPM)_adj_ change was in favor of the same experimental condition as the change in ΔΔlog_2_(TPM)_norm_, then the segment use was considered to be the location of a specific usage change (“changepoint”). If the P value of the change in ΔΔlog_2_(TPM)_adj_ was not significant, then the difference in usage was simply marked in favor of the condition with greater usage of that segment. In the case of the last segment (the end of the 3’UTR), if ΔΔlog_2_(TPM)_norm_ was not significant between conditions, the abundance of that segment between conditions was considered identical. If ΔΔlog_2_(TPM)_norm_ was significant between conditions, the abundance of that segment between conditions was marked as higher in one condition over the other. In this way, every segment of the mRNA was quantified between conditions: the first segment was quantified based on log_2_FC of TPM between experimental conditions, and every other segment was quantified based on whether it is differentially used between the two conditions.

Each of these segment-by-segment comparisons was then given a score. For convenience, one condition was assigned negative values while the other was assigned positive values. Usage differences of the first segment were arbitrarily awarded a score of 3 or −3 depending on which condition had the higher abundance. Internal segments were given values of 1 or −1 depending on which condition used them more. In cases where there was no difference in relative use, the score is 0. However, when an internal segment was considered a changepoint, that segment was awarded a 3 or −3. Last segments were awarded 4 or −4 depending on condition, thus the weight of the last segment was the highest. Where there was no difference in the last segment, the scores of all the internal segments were summed, and the score of the last segment was −4 if the sum was negative, 4 if the sum was positive, and 0 if the sum was 0.

Finally, the scores of the first segment and the last segment for each transcript were summed in order to determine whether the transcript was longer in one condition or the other. Scores of +/-1, +/-4, +/-7 indicate that the transcript tends have a longer 3’ UTR in one condition over the other, while scores of 0, 3, −3 indicate that a transcript undergoes no APA although there may be an abundance difference of the transcript as a whole between conditions. It is for this reason that the difference in last segment usage in Figure 2A may not be significant but the ultimate decision on whether the longer 3’UTR is differentially used between conditions falls in favor of one condition or the other.

### Transcript-level changes in abundance

For transcript-level analyses, kallisto was used for the pseudoalignment of RNA-Seq reads against the standard RefSeq transcriptome (Bray et al., 2016). Abundance changes between conditions were calculated by averaging the TPMs for each transcript across trials and then calculating the log_2_FC between conditions. kallisto was used for this process in order to keep the tools of this study consistent, but the results of this analysis conformed to the outcome of DESeq2 analysis on the same dataset (data not shown; (Love et al., 2014).

### Metabolic labeling

The method used for metabolic labeling has been documented in our previous work (Kishor et al 2019, Baird et al 2018). Briefly, HEK-293TO cells were reverse-transfected with a gene-specific or non-targeting control siRNA for 72 h. At the end of the depletion, cells were treated with 0.5 mM 5-ethynyl uridine (5-EU) for 60 min. RNA was isolated using TRIzol. Total and nascent RNA levels in each sample was partitioned using the Click-iT Nascent RNA Capture Kit (ThermoFisher Scientific, Cat. No. C10365) following the manufacturer’s protocol. mRNA abundance was determined using qRT–PCR. Individual half-lives were determined using the equation: *t*1/2 *=* −*tL \** ln(2)/ln(1/R), where *tL* is the 5′EU labeling time in minutes and *R* is the abundance in nascent RNA fraction/abundance in total RNA fraction (Haque et al., 2018; Russo et al., 2017). At least 4 independent biological replicates were performed for each experimental condition.

### UPF1 RNA affinity purification

HEK-293 FLP-In stable cell lines expressing CLIP-tagged UPF1 (above) were seeded in 6 x 15 cm plates and then treated with 200 ng/mL doxycycline hyclate (Sigma) for 48 hours to induce CLIP-UPF1 expression. In parallel, a human cell line that had been stably integrated with GFP was used as a control. Cells were harvested 48 hours post-induction at 80-85% confluency and whole cell lysate generated by the freeze/thaw method as previously described (Fritz et al., 2018; Hogg and Collins, 2007a, 2007b; Hogg and Goff, 2010). The cell extracts were equilibrated to 2.7 mg/mL with HLB-150 (20 mM HEPES-NaOH pH 7.6, 150 mM NaCl, 2 mM MgCl_2_, 10% glycerol) supplemented with 1 mM DTT and 0.1% NP-40 (final) to achieve a final volume of 1 mL. Lysates were spun at top speed (18,000 *g*) in a microcentrifuge pre-chilled to 4°C, and 800 µL was transferred to a 1.7 mL microcentrifuge tube. The remaining cell extract was reserved for downstream input analysis by Western blot (2.5 µL) and high-throughput sequencing plus RT-qPCR (100 µL). The total volume was brought to 900 µL by the addition of 91 µL of HLB-150 supplemented with 1 mM DTT and 0.1% NP-40 and 9 µL of 1 mM CLIP-Biotin (New England Biolabs; 10 µM final). The samples were rotated (end-over-end) for 1 hour at 4°C to allow CLIP-UPF1 to react with CLIP-Biotin substrate to yield covalently biotinylated protein. Unbound CLIP-Biotin was subsequently removed by passing the samples (750 µL) through Zeba™ Spin Desalting Columns, 40K MWCO, 2 mL (Thermo Scientific) according to manufacturer instructions. Buffer exchange was performed with HLB-150 supplemented with 1 mM DTT and 0.1% NP-40. 240 µL of pre-washed Dynabeads^TM^ MyOne^TM^ Streptavidin T1 (Invitrogen) was then added and the samples rotated (end-over-end) for 1 hour at 4°C to immobilize the biotin-bound CLIP-UPF1 complexes. The samples were then washed three times with 500 µL of HLB-150 supplemented with 1 mM DTT and 0.1% NP-40 and one one-hundredth reserved for downstream Western blot analysis of CLIP-UPF1 pull-down efficiency. The remainder was combined with 250 µL TRIzol^TM^ (Invitrogen) and RNA was isolated according to manufacturer instructions. DNase-treatment was subsequently performed using RQ1 RNase-Free DNase (Promega) and RNA isolated by acid phenol chloroform extraction according to standard protocol. This resulted in approximately 1 µg of total RNA that was then subjected to high-throughput sequencing. In parallel, the reserved 10% of equilibrated lysate (approximately 3 µg of total RNA) was also sent for high-throughput sequencing. A total of three biological replicates from each condition were processed. Sequencing libraries were prepared from input and bound RNA using the Illumina TruSeq Stranded Total RNA Human kit and sequenced on an Illumina HiSeq 3000 instrument.

### qRT-PCR

For steady state samples, RNA was extracted from cells using the TRIzol reagent and treated with RQ1 DNAse (Promega, Madison, WI). 500 ng of each sample was used for cDNA synthesis using the Maxima First Strand cDNA Synthesis Kit for RT–qPCR (Thermo Scientific, Philadelphia, PA). cDNAs were diluted 1:20 with water and used for qPCR with iTaq Universal SYBR Green Supermix (Bio-Rad, Hercules, CA) on a LightCycler 96 thermocycler (Roche, Basel, Switzerland). For steady-state relative isoform abundance quantification, abundances were calculated by the ΔΔCt method normalized to the abundance of the CDS primer pair and then the non-targeting condition. At least 3 independent biological replicates were performed for each condition, and statistical significance was assessed by two-tailed Student’s t-test. For metabolic labeling samples, reverse transcription was carried out as part of the Click-iT Nascent RNA protocol and qPCR was completed as above for nascent and total RNA fractions. Abundances were calculated by the ΔΔCt method normalized to the abundance of GAPDH. The cDNA abundance in each biological replicate was assessed at least twice to minimize technical variation. Primer sequences are listed in Table S4.

### Software

Analysis of pA segments was performed using JMP 14.0.0. Graphical representation of RNA-Seq reads and eCLIP peaks was generated by the Integrative Genomics Viewer (IGV 2.3.68). Statistical tests and half-life calculations were performed using Microsoft Excel version 16.28.

