## Supplemental Information for "Activation and inhibition of nonsense-mediated mRNA decay controls the abundance of alternative polyadenylation products"

**Supplemental Tables**

**Table S1. Segment-level analysis of RNA-Seq of cells treated with siUPF1 or siNS.**

**Table S2. Segment-level analysis of UPF1 RIP-Seq.**

**Table S3. Segment-level analysis of RNA-Seq of cells treated with siPTBP1, siUPF1, siPTBP1/siUPF1, or siNS.**

**Table S4. Primers used in this study.**

### Figure S1

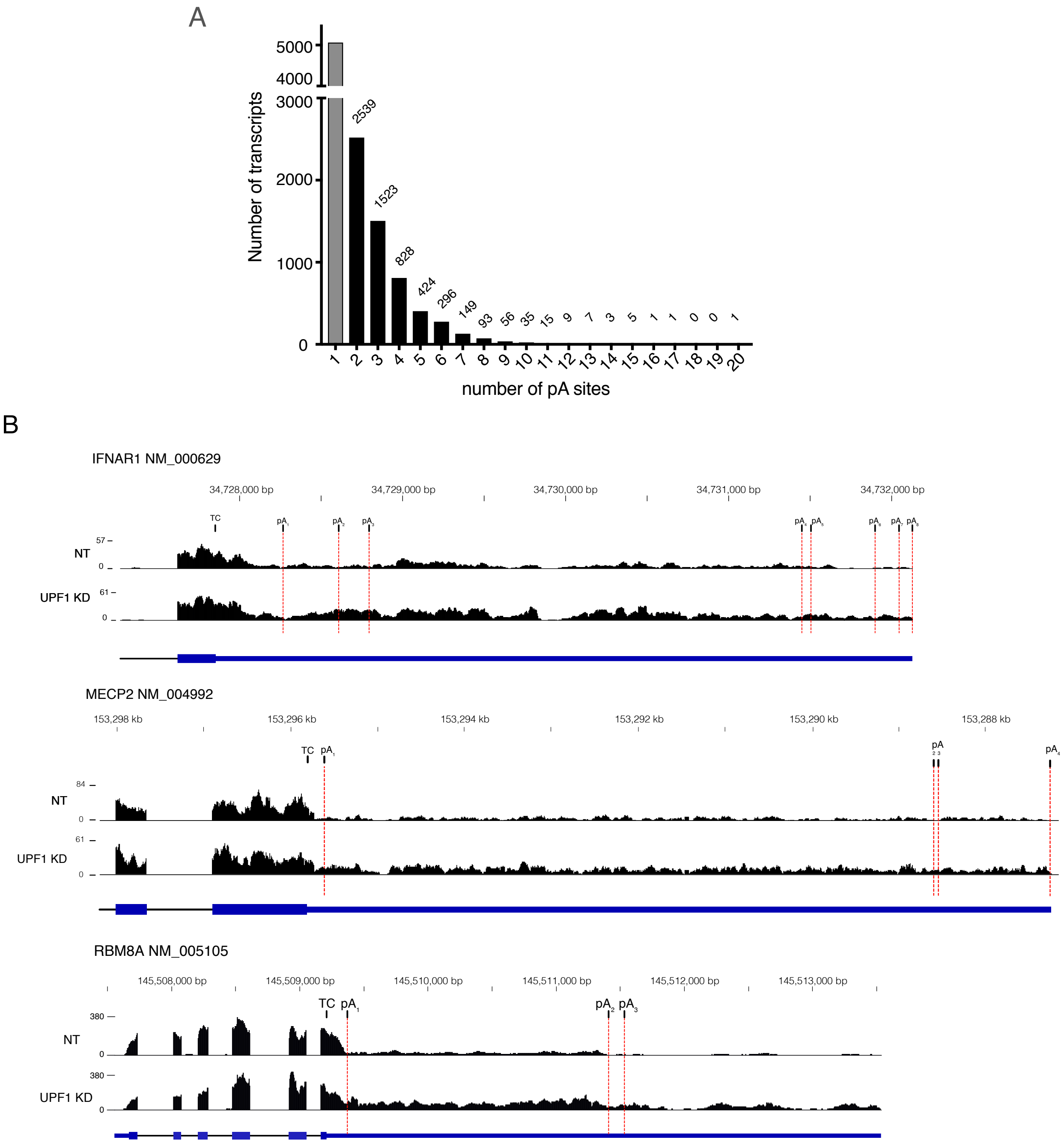


**Figure S1. Segment-level analysis of APA product abundance. (A)** Histogram of the number of polyA sites in the custom transcriptome. Transcripts with only 1 polyA site are in grey since they are not included in the analyses of distal usage. The total number of transcripts in each bin is indicated above each bar. **(B)** Example traces of transcripts identified as having increased long 3'UTR isoform expression upon UPF1 KD. The location of pA sites is indicated, and red lines indicate boundaries of segments used for quantification.

### Figure S2

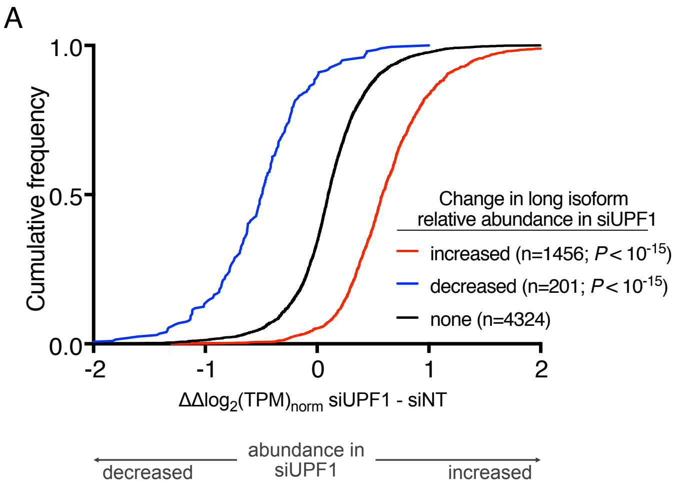


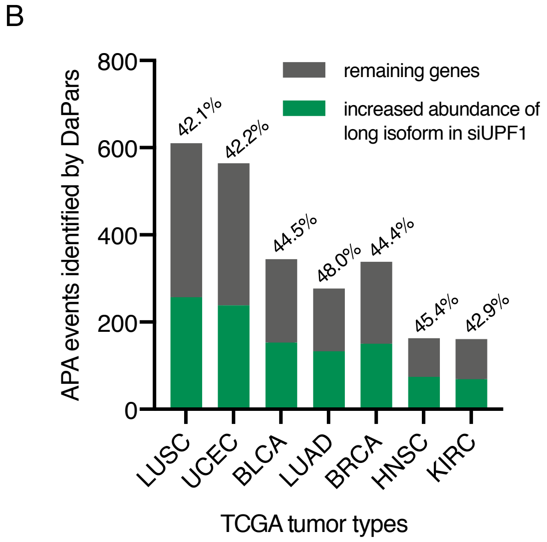


C

Genes with APA in tumors and long isoform NMD sensitivity

| Identified in 6 tumor types | | Identified in 7 tumor types |
|---|---|---|
| PNPT1 | TMEM167A | RPS10-NUDT3 |
| CDS2 | CSNK2A1 | TMEM237 |
| CEP104 | YIPF6 | SRI |
| SPTLC2 | CHURC1 | MRS2 |
| CSNK1D | FLNB | CIAO1 |
| BID | COMMD2 | ASB1 |
| PPP6R2 | TFDP2 | SAP30L |
| LANCL2 | ESYT2 | C6orf89 |
| SLC25A37 | SYNCRIP | UBR2 |
| DPH3 | | MECP2 |
| PA2G4 | | PPP1CB |
| SH3GLB1 | | ATL3 |
| PAFAH1B1 | | TMEM106B |
| HSPA4 | | ABI2 |
| ARIH2 | | TMEM64 |

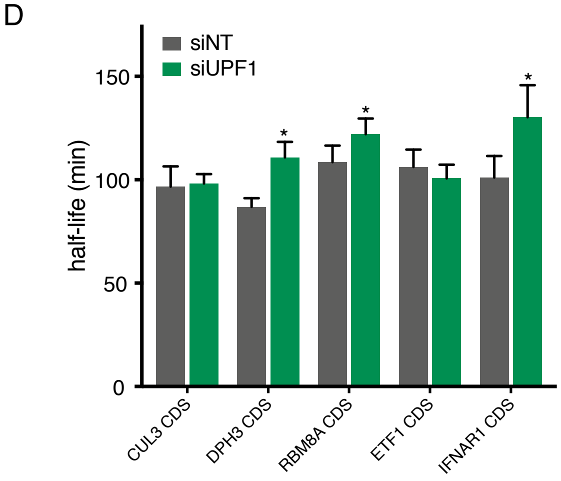


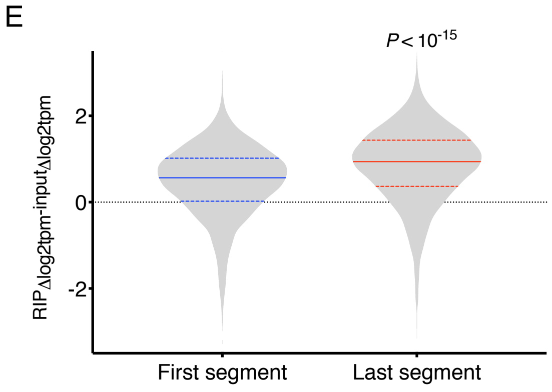


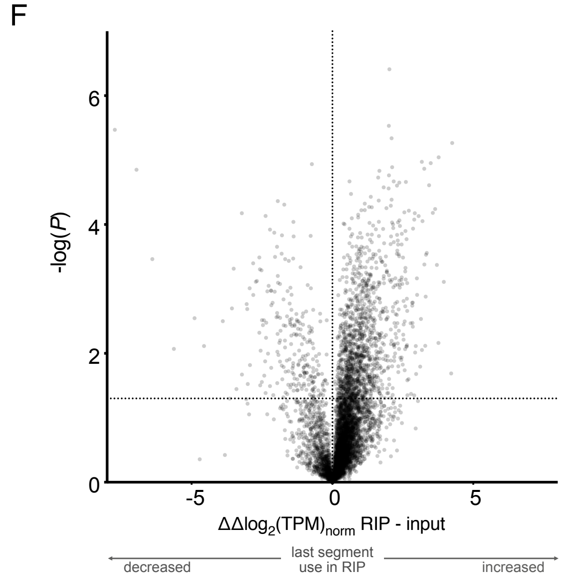

**Figure S2. Characteristics of transcripts bound and regulated by UPF1. (A)** CDF plot representing $\Delta\Delta log_2$(TPM)norm between non-targeting and UPF1 knockdown of the last segment. Traces represent classes of transcripts from the results of the decision tree. Significance was calculated using the K-S test. **(B)** Proportion of transcripts with increased long isoforms upon UPF1 knockdown in the set of APA events in common with each of the 7 tumor types (LUSC, lung squamous cell carcinoma; UCEC, uterine corpus endometrial carcinoma; BLCA, bladder urothelial carcinoma; LUAD, lung adenocarcinoma; BRCA, breast invasive carcinoma; HNSC, head-neck squamous cell carcinoma; KIRC, kidney renal clear cell carcinoma) evaluated by DaPars (Xia et al., 2014). **(C)** Genes identified as undergoing increased long 3'UTR isoform expression upon UPF1 knockdown and APA in six or seven of the tumor types in B. Genes confirmed by qRT-PCR in this study are underlined. **(D)** Metabolic labeling was used to determine the half-lives of all isoforms containing the coding sequence under conditions of UPF1 knockdown or non-targeting siRNA (n =4). Significance was determined using a two-tailed Student's t-test. For clarity, a single star indicates $P \leq 0.05$ when compared to the non-targeting condition. Error bars represent 1 standard deviation. **(E)** Distribution of the abundance change between UPF1-associated RNA and the input RNA for first segments and last segments of transcripts from the segment-level analysis. Significance was calculated using the K-S test. **(F)** Volcano plot representing the $\Delta\Delta log_2$(TPM)norm between affinity purified and input RNA for transcripts of the last segment. Horizontal line indicates the significance threshold equal to $P \leq 0.05$ (n = 3).

### Figure S3

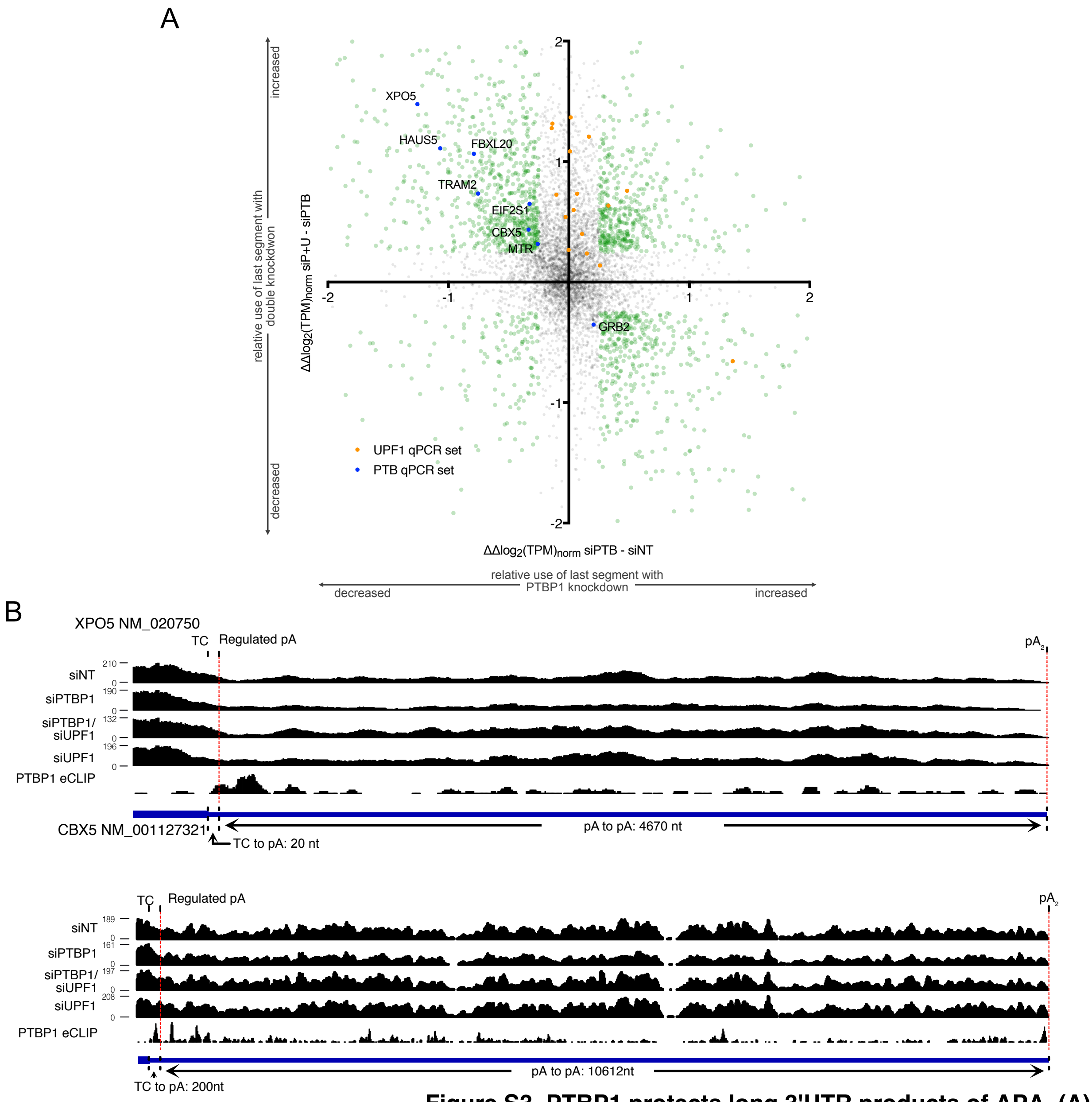


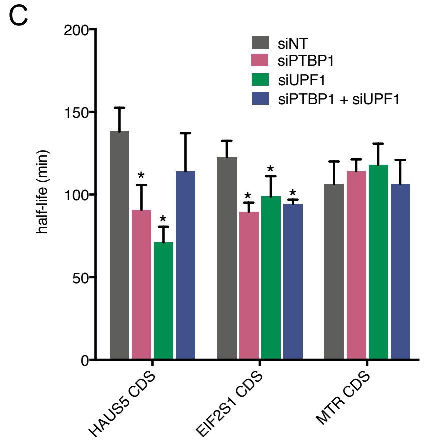


**Figure S3. PTBP1 protects long 3'UTR products of APA. (A)** Scatter plot representing the last segment $\Delta\Delta log_2$(TPM)norm between non-targeting and PTBP1 knockdown for transcripts with more than 1 polyA site against the $\Delta\Delta log_2$(TPM)norm between PTBP1 + UPF1 KD and PTBP1 KD. Gray points are transcripts that are below the threshold for significant change. Orange points are the transcripts that were qPCR validated as part of the UPF1 KD analysis and blue points are those that were qRT-PCR validated as part of the PTBP1 analysis. **(B)** Additional example traces of transcripts identified by segment-level analysis as having long 3' UTR isoform abundance restored by UPF1 + PTBP1 knockdown. The location of pA sites is indicated, and red lines bound segments used for quantification. **(C)** Metabolic labeling to determine the half-lives of the transcript CDS under the indicated conditions (n=4). Significance was determined using a two-tailed Student's t-test. (*$P \le 0.05$; error bars represent 1 SD).
